# Aspects of the glycome of endochondral ossification

**DOI:** 10.1101/2022.05.12.491629

**Authors:** Sheena F McClure, John McClure

## Abstract

Comparatively little is known about the glycome (the set of glycans and glycoconjugates made by the cell, tissue or organism) of bone and cartilage. The glycome has a high-density coding capacity and is important in post-translational modification. Lectin histochemistry provides insights into the glycome and we applied this technique to an ectopic ossification site in human bronchial cartilages.

Our results show that cartilage matrix at the site of erosion by chondroclasts has a limited but definite glycoprofile. In contrast, bone matrix does not express many lectin ligands. Chondroclasts have an extensive glycoprofile similar to that of, but not identical to, osteoclasts. N and O glycans are both expressed in the zone of presumptive mineralization and are relevant to that process. Bone trabecular lining cells communicate with osteocytes via intracanalicular processes and some lectin ligands were expressed by these three components.

Mast cells and angiogenesis were prominent. Since cartilage normally resists vascular penetration by the secretion of antiangiogenesis factors it is postulated that the hypertrophic chondrocytes in the mineralization zone produce proteases which inhibit antiangiogenesis and facilitate angiogenesis by mast cells.

## INTRODUCTION

The glycome is the complete set of glycans and glycoconjugates made by a cell, tissue or organism (1). These molecules form a ‘sugar code’ which has fundamentally changed understanding of the significance of glycosylation. They have a high-density coding capacity and are active in post-translational modifications (2).

Despite its evident importance very little is known about the glycome of bone tissue. Somewhat more is known about human articular cartilage and its related sub-chondral bone plate (3,4). In embryonic development the skeleton is initially modelled in cartilage to be replaced by bone, except at the epiphyseal plates, by which postnatal growth to maturity is achieved. The cartilage replacement, both initially and part of the growth plate activity, is by a process of endochondral ossification. There is some limited glycomic information on this but it is restricted to animal studies. Outside the skeleton, cartilage is present in the trachea and major bronchi. We noted that in some of these ectopic cartilages there may be partial replacement with bone tissue by an endochondral ossification to produce complete ossicles. We examined these by lectin histochemistry which provides information about the composition of the glycome (5).

The results form the basis of this report.

## MATERIALS AND METHODS

In a lectin histochemical study of lung, 19 formalin-fixed, paraffin wax-embedded blocks were retrieved from a histopathology archive. The blocks contained bronchial tissue with cartilage showing partial replacement by bone tissue. After decalcification tissue sections were stained with a panel of 10 biotinylated lectins (Table 1) using our standard lectin histochemical techniques with appropriate controls (6).

**TABLE 1.**

| LECTIN | ORIGIN/SOURCE | MAJOR SPECIFICITY |
| --- | --- | --- |
| HHA | Hippeastrum hybrid (amaryllis) | α1,2,α1,3 and α1,6 - Mannose |
| PSA | Pisum sativum (garden pea) | α-D-Mannose in non-bisected bi/tri antennary, complex N-glycans |
| ePHA | Phaseolus vulgaris (kidney bean) | Bi/tri antennary, bisected complex N-glycans |
| IPHA | Phaseolus vulgaris (kidney bean) | Tri/tetra antennary, non-bisected complex N-glycans |
| AHA | Arachis hypogaea (peanut) | Gal1,3GalNAcα1>Galβ1,4GlcNAcβ1 |
| HPA | Helix pomatia (Roman snail) | GalNAcα1 |
| MPA | Maclura pomifera (osage orange) | Galβ1,3GalNAcα1>GalNAcα1 |
| ECA | Erythrina cristagalli (coral tree) | Galβ1,4GlcNAcβ1 |
| SBA | Glycine max (soy bean) | Terminal GalNAcα1Galα1 |
| SWGA | Triticum vulgaris (wheat germ) | (Galβ1,4GlcNAcβ1)<sub>n</sub> and GalNAc |

Origin and carbohydrate specificities of lectins used in this study.

## RESULTS

Partial ossification had arisen by a process of endochondral ossification creating an ossicle with external cortical type bone and internal trabecular and intertrabecular tissues. Bone structures stained with 10 lectins (HHA, PSA, ePHA, lPHA, sWGA, AHA, ECA, MPA, HPA and SBA). The osteoblast/trabecular lining cell layer stained with HHA, PSA, ePHA, sWGA and SBA accounting for 23% of reactions. Osteocytes stained with HHA, PSA, ePHA, sWGA and SBA accounting for 38.5% of reactions. Osteoclasts stained with HHA, PSA, lPHA, sWGA, AHA, ECA, HPA, MPA and SBA also accounting for 38.5% of reactions. The cartilage matrix at the sites of endochondral ossification stained with ePHA, lPHA, AHA and SBA. Erosion of this matrix was by chondroclasts which stained with ePHA, 1PHA, sWGA, AHA and SBA. There was ensuing bone deposition. Mast cells in the intertrabecular spaces stained with HHA, PSA, ePHA, lPHA and adjacent blood vessels with HHA, lPHA, AHA, ECA and SBA.

In the instances where the trabecular lining cell layer and osteocytes stained with the same lectin, canalicular communications between the two also stained.

In general terms the observed staining reactions were cytoplasmic and membrane sited. Nuclear staining was never observed.

In contrast to the cartilage matrix in the zone of endochondral ossification, the bone matrix generally did not stain with lectins. There was occasional very faint matrical staining by ePHA, lPHA, sWGA and SBA.

## DISCUSSION

The lectins used and their specificities are given in Table 1. There is very little published information on the glycome of bone and its constituent structures either in health or disease. Osteoclasts have been shown to have ligands for AHA (PNA) and WGA (7,8). Lyons et al (4) have published data on the subchondral bone plate of the normal human knee.

In the present study there were staining reactions for osteoblasts/trabecular lining cells, osteocytes and osteoclasts. The particular reactions of these and chondroclasts are shown in Table 2.

**TABLE 2.**

| LECTIN | OSTEOBLASTS/TRABECULAR LINING CELLS | OSTEOCYTES | OSTEOCLASTS | CHONDROCLASTS |
| --- | --- | --- | --- | --- |
| HHA (N) | + | + | + | - |
| PSA (NC) | + | + | + | - |
| ePHA (NC) | + | + | - | + |
| IPHA (NC) | - | + | + | + |
| AHA (O) | - | + | + | + |
| HPA (O) | - | - | + | - |
| MPA (O) | - | + | + | - |
| ECA (O) | + | + | + | + |
| SBA (O) | - | - | - | + |
| SWGA (O) | + | + | + | + |

(N = N-glycans; NC = Complex N-glycans; O = O-glycans; + = reaction; - = no reaction).

Overall the observed glycomes are limited with the osteoblast/trabecular lining and chondroclasts more so than osteocytes or osteoclasts.

Osteoblasts are actively bone-forming cells with a typical cubical shape. When inactive they become flattened onto the trabecular surfaces forming the majority of the trabecular lining cell layer. Where the staining reactions (HHA, PSA, ePHA, SBA, sWGA) are common to the trabecular lining cell and osteocyte there was staining of the osteocytic intra-canalicular processes communicating with the surface layer. Osteoclasts and osteocytes have similar but not quite identical repertoires of lectin ligands and both are more extensive than that of osteoblast/trabecular lining cells. All three stained with HHA indicating N-glycans binding. Similarly with PSA showing N-glycan binding enhanced by core fucosylation. Ligands for SBA (terminal GAlNAcα1 > Galα1) and sWGA ((3Galβ1,4GlcNAcβ1-)_n_ and GalNAc) were common to the three cell types confirming O-glycosylation. There were bisected complex-type N-glycans in osteoblast/trabecular lining cells and osteocytes (ePHA) and tetra- and tri-antennary N-glycans in osteocytes and osteoclasts (lPHA). Uniquely osteoclasts expressed the ligand for HPA (GalNAcα1-).

The mast cell glycome was limited to four interactions. HHA staining is a common feature of several reports on mast cell ligand expression (9). The presence of complex N-glycans is demonstrated by PSA, ePHA and lPHA.

Blood vessels not only contained complex N-glycans but also O-glycans and O-glycosylation is a reported feature of endothelial cells (10).

The staining of osteocytes, osteoclasts, mast cells and blood vessels with lPHA is of particular significance. The specific ligand for this lectin is the β1-6 branching linkage in complex N-glycans and it is formed by the action of GnTaseV (MGAT5). This enzyme is overexpressed by carcinomas and is believed to promote tumour growth and metastasis probably via angiogenesis (11). Mast cells also promote angiogenesis and the expression of lPHA ligand by osteoclasts, mast cells and blood vessels may represent a link between bone resorption and the accompanying neovascularisation.

The ossicles had arisen by an endochondral ossification. On the external aspect there were foci of cartilage undergoing resorption by chondroclasts and replacement by bone. The constituent chondrocytes were hypertrophic and their matrix stained with PSA, ePHA, lPHA, AHA and sWGA indicating complex N-glycans and O-glycosylation. Although the chondroclasts were morphologically identical to osteoclasts, they had a more limited glycoprofile and did not stain with HHA, PSA, HPA, MPA or ECA. Thus there were ligands for complex N-glycans (ePHA, lPHA) and for O-glycans (AHA, SBA, sWGA). The expression of lPHA ligand may again be related to angiogenesis.

The endochondral ossification observed here shares the features of cartilage calcification, its erosion and replacement by bone, with those in the growth plate. However, it is less complex architecturally and easier to examine in tissue sections. Also it is neither weight-bearing nor part of an articulation. Calcification of cartilage is a key process in endochondral ossification and there is evidence that the presence of N- and O-glycans may be relevant to mineralization.

O-glycosylation is initiated by the transfer of an N-acetylgalactosamine residue (GalNAc) to the hydroxyl group of either serine or threonine to form Tn antigen (GalNAcα1-S/T). This and further modifications are controlled by a group of enzymes (UDP-N-acetylgalactosamine: polypeptide N-acetylgalactosaminyltransferases ppGalNAcTs or ppGalNTases).

In familial tumoural calcinosis there is hyperphosphataemia and ectopic calcified masses and some families have mutations in the gene for the glycosyltransferase ppGalNAc-T3. Other families have mutations in the phosphate-regulating Fibroblast Growth Factor 23 (FGF23) indicating a possible link between FGF23 and O-glycosylation. FGF23 negatively regulates the resorption of phosphate from the renal proximal tubule. Absent or defective FGF23 activity will result in hyperphosphataemia. Patients with mutations in ppGalNAc-T3 have increased levels of inactive and low levels of active FGF23. It is believed that O-glycans normally present on FGF23 confers protection from proteolysis and aids the production of intact FGF23 (12).

Certain mesenchymal tumours produce FGF23 resulting in hyperphosphaturia, hypophosphataemia and oncogenic osteomalacia (13).

Another line of evidence is provided by reported studies on immortalized human mesenchymal stromal cells (MSC). Inhibiting O-glycan processing in the Golgi apparatus prior to the start of osteogenesis inhibits the mineralization capacity of the later formed osteoblasts. In contrast, inhibiting the N-glycan processing enhances the mineralization capacity of the osteoblasts (14).

FGF23 as indicated above has a major role in phosphate homeostasis. However, it is also an inhibitor of mineralisation although it is unclear if this is a direct or indirect effect (15). Originally located in the ventrolateral nucleus of the brain (16) it is now known to be mainly secreted by skeletal osteocytes (17). FGF23 overexpression supresses not only osteoblast differentiation but also matrix mineralization independently of its systemic effects on phosphate homeostasis (18,19). Shalhoub et al (20) also noted inhibition of mineralization but found that FGF23 and its cofactor alpha Klotho in excess stimulated osteoblastic MC3T3.E1 cell proliferation.

These data have been obtained from animal models and in vitro experiments, refer to cells of an osteoblastic lineage (with no mention of a chondroblastic one), and are likely to be context specific. If, or how, these results transfer to the human skeleton is undetermined. However, the osteocyte, which at one time was considered to be inert, is emerging as a major player in skeletal homeostasis, in addition to its role as a mechanosensory cell. In the adult skeleton the osteocyte forms 90-95% of the bone cell population which in addition to its endocrine function in phosphate metabolism is likely to have other autocrine and paracrine functions (21). Perhaps local control of mineralization is one of these.

The cellular controller of the local mineralization of cartilage matrix is the hypertrophic chondrocyte which produces collagen type X, matrix vesicles and alkaline phosphatase (22,23). This cell features the expression of both N-glycans and O-glycans ligands (24) and the question arises as to their roles in the present study. Collagen type X has a highly conserved N-glycosylation site within its C-terminal and its function has remained elusive. Recently, it has been shown that whilst dispensable under normal conditions N-glycans are essential for collagen folding and secretion under conditions that challenge proteostasis such as during development, tissue repair and disease (25).

Matrix vesicles are small packages released from the hypertrophic chondrocyte and which mineralize the matrix by associating with collagen type X. They require calcium and phosphate ions. The latter can be released by alkaline phosphatase from appropriate substates but FGF23 and Klotho may also contribute and O-glycosylation is needed to protect FGF23 from premature proteolysis. In the proximal renal tubular cell FGF23 reduces the expression of type II A and C sodium-dependent phosphate co-transporters (NPT2 A & C) leading to decreased cellular phosphate resorption and raising the phosphate levels in the renal interstitium (19). Perhaps there is a similar effect on the hypertrophic chondrocyte.

Like osteoclasts, chondroclasts can only resorb fully mineralized surfaces and cartilage will normally resist resorption until mineralization occurs (26).

Cartilage also normally resists vascular penetration by the secretion of antiangiogenesis factors. Hypertrophic chondrocytes produce proteases which inhibit antiangiogenesis and facilitate angiogenic factors possibly produced by the observed mast cells. This combination of matrix calcification and angiogenesis permits replacement of previously resistant cartilage by bone (27).

## Figures

**Figure A.**
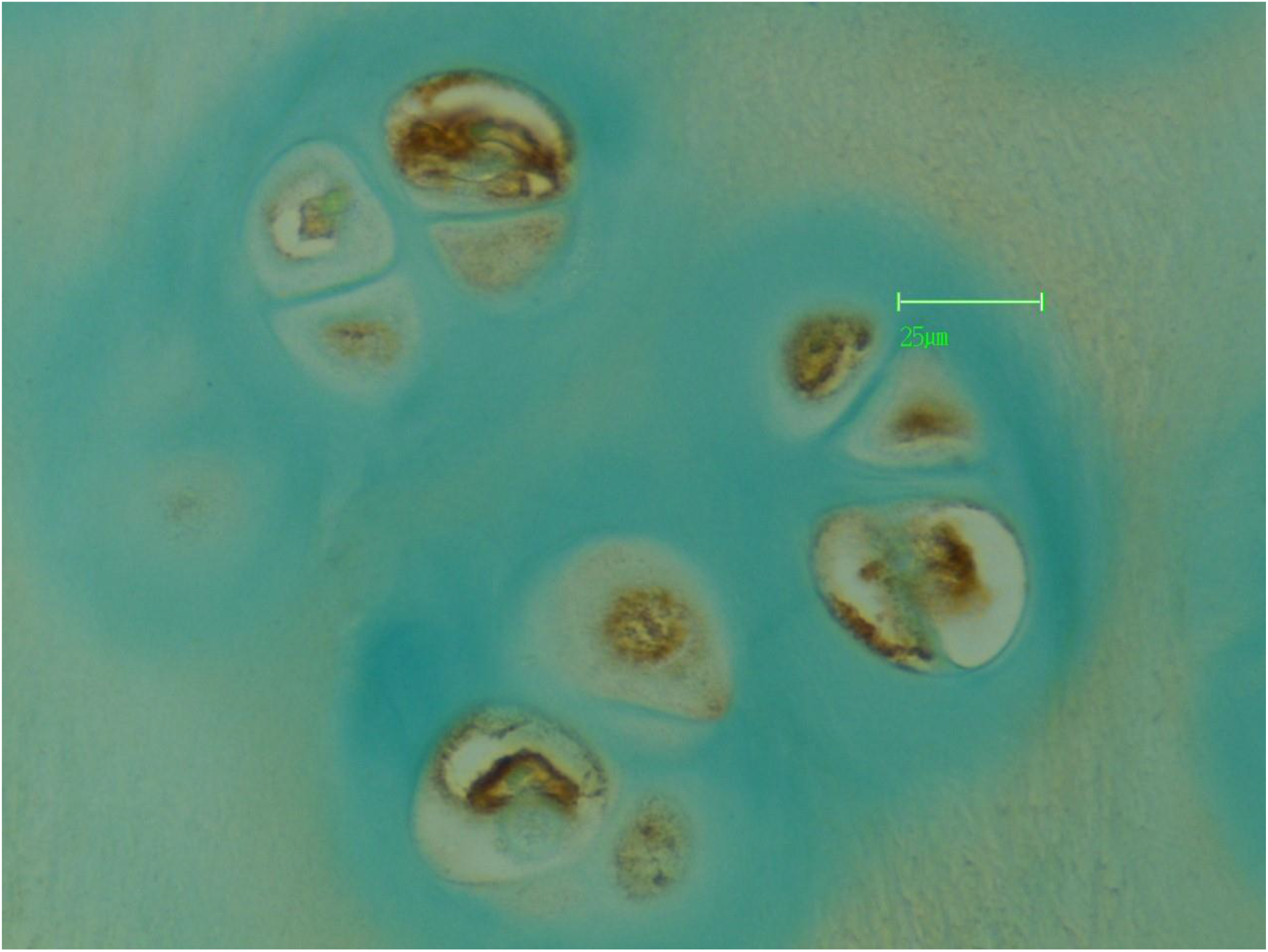
The hypertrophic cartilage cells show positive staining of cytoplasm and cell membranes by PSA lectin indicating the presence of complex N-glycans.

**Figure B.**
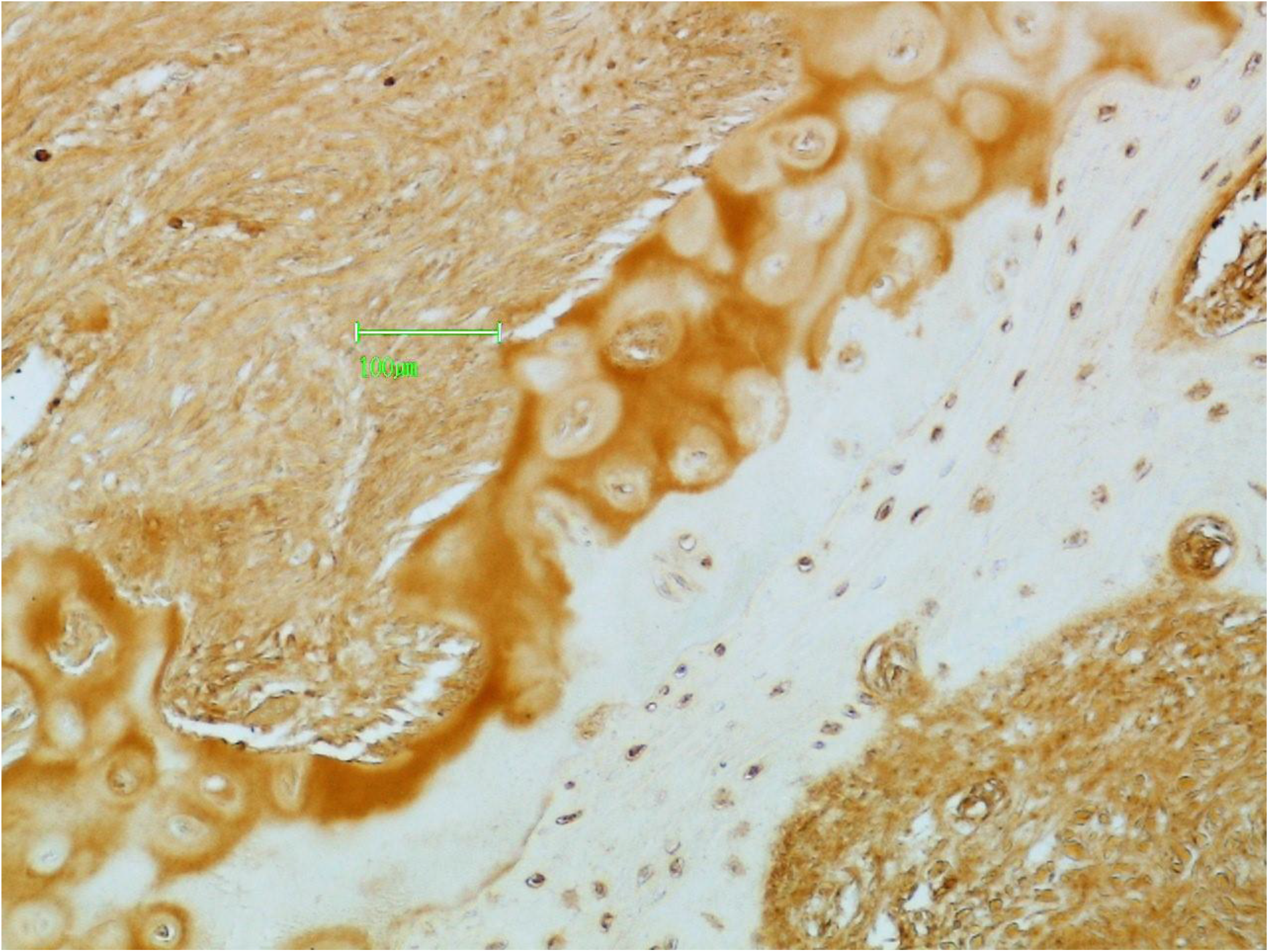
The matrix of the hypertrophic cell zone is also stained by PSA lectin again indicating the presence of complex N-glycans.

**Figure C.**
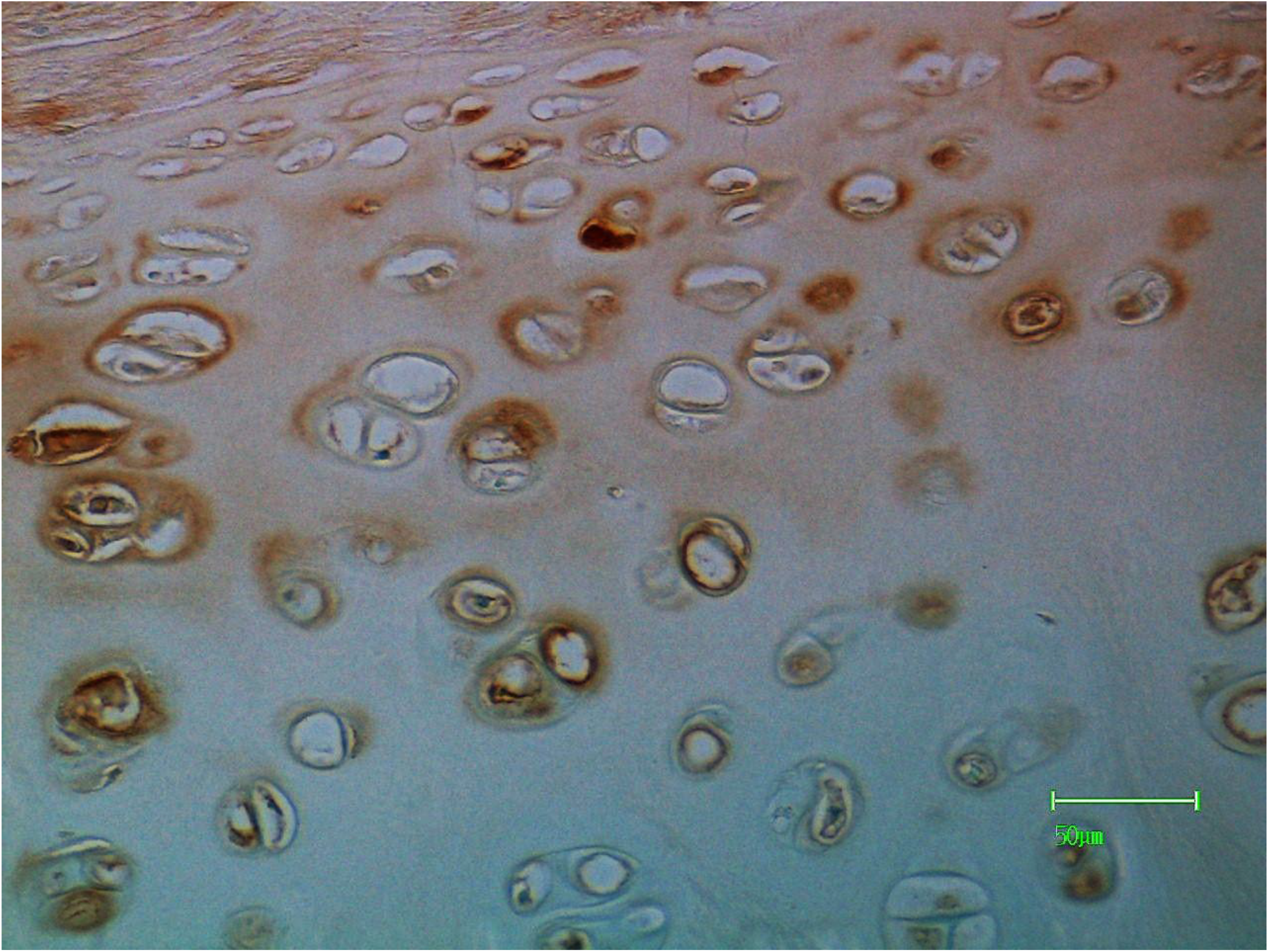
The cytoplasm, membrane and adjacent matrix of hypertrophic cartilage cells are stained by AHA lectin indicating the presence of O-glycans.

**Figure D.**
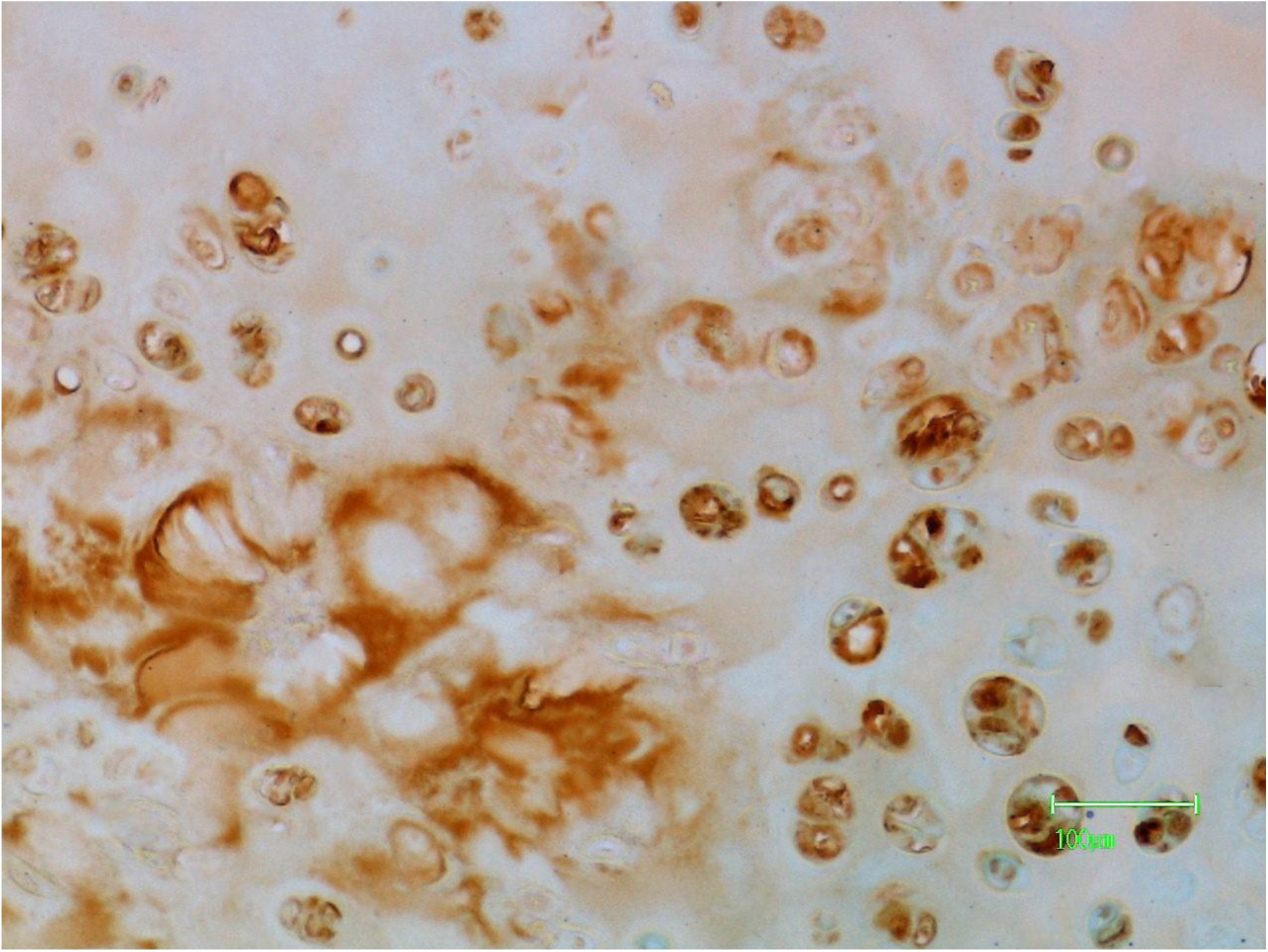
The hypertrophic cartilage cells and matrix are stained by sWGA lectin indicating the presence of O-glycans.

**Figure E.**
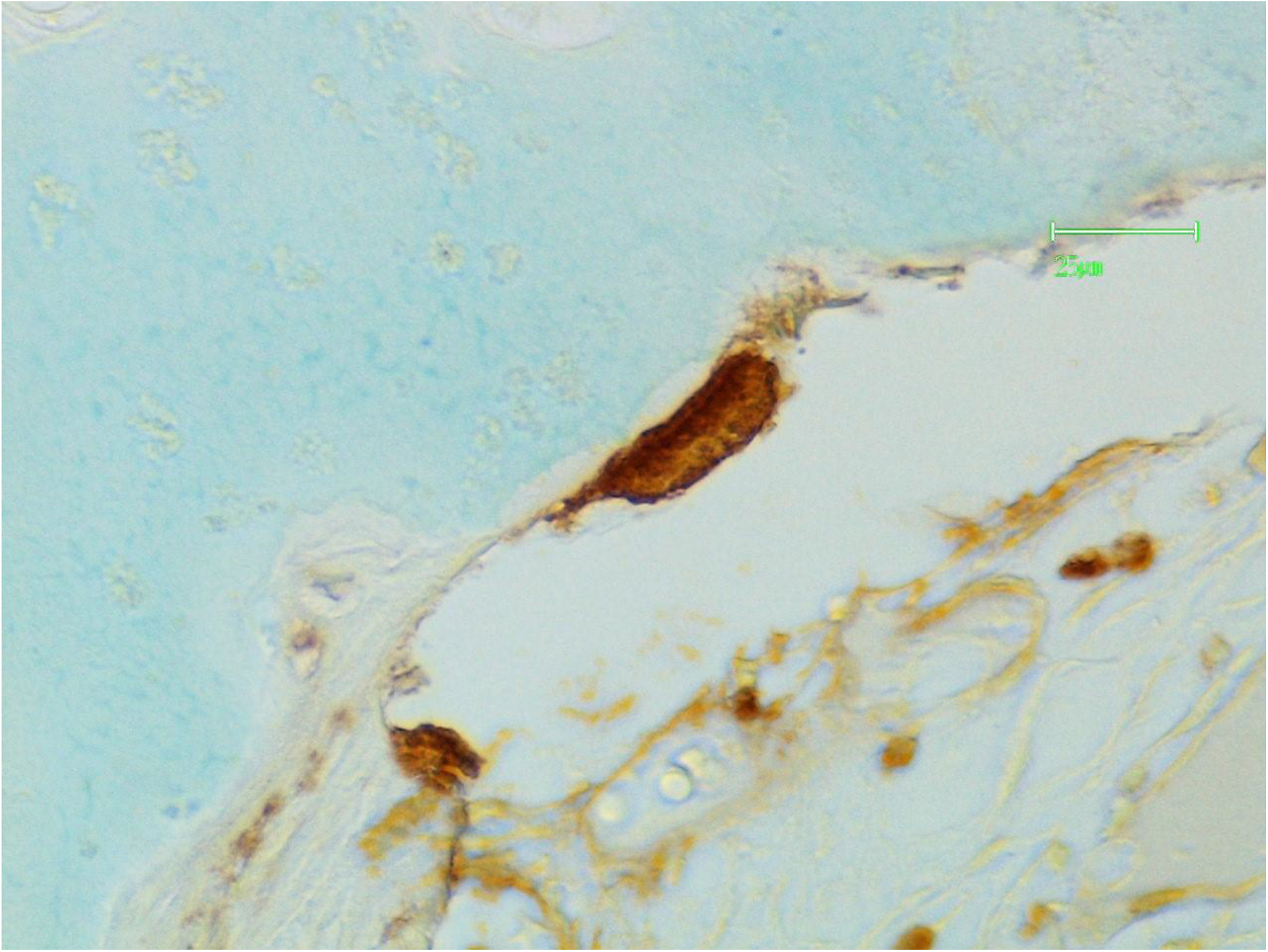
The chondroclast is stained by lPHA lectin indicating the presence of complex N-glycans.

**Figure F.**
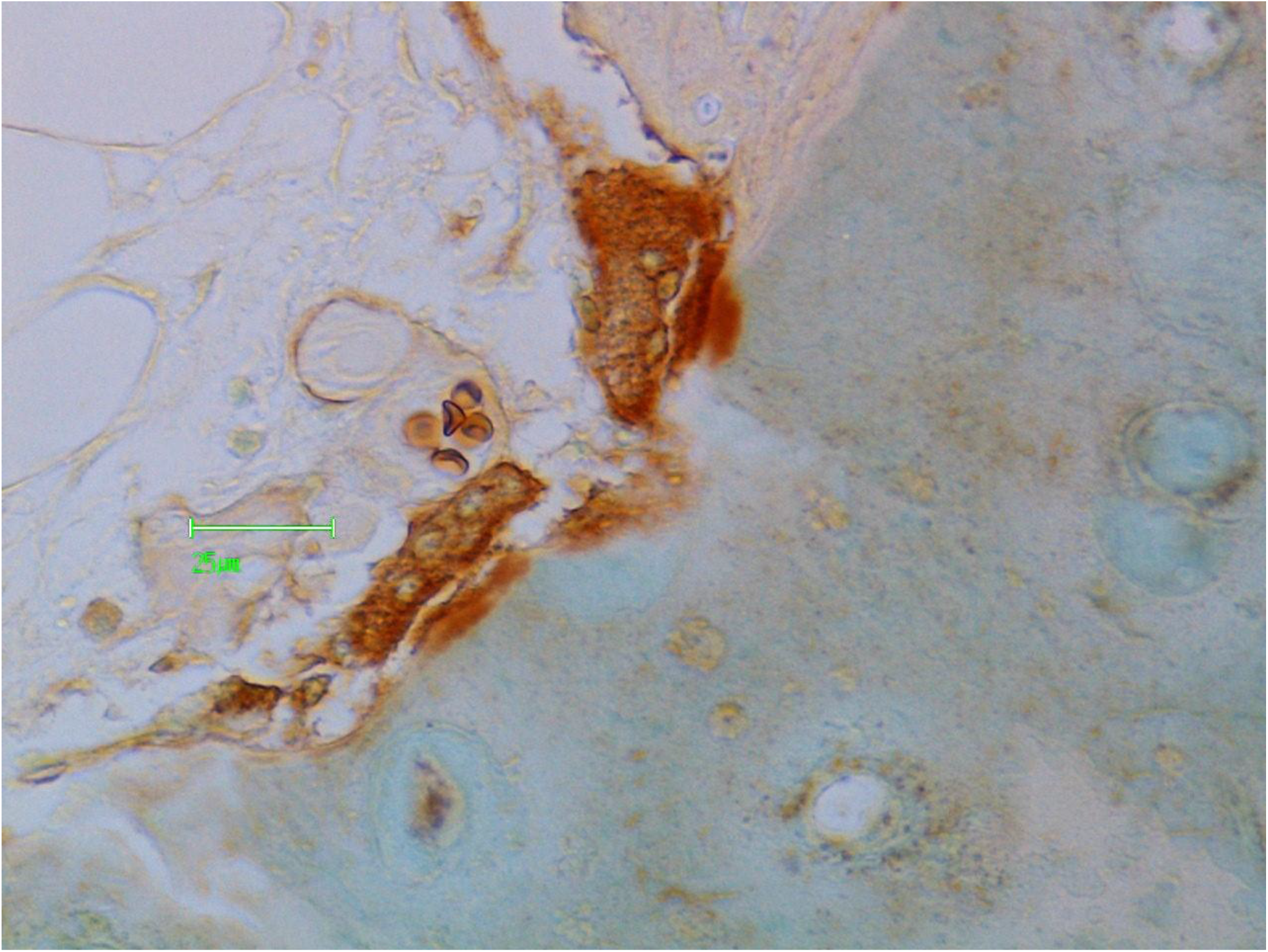
Chondroclasts are stained by sWGA lectin indicating the presence of O-glycans.

**Figure G.**
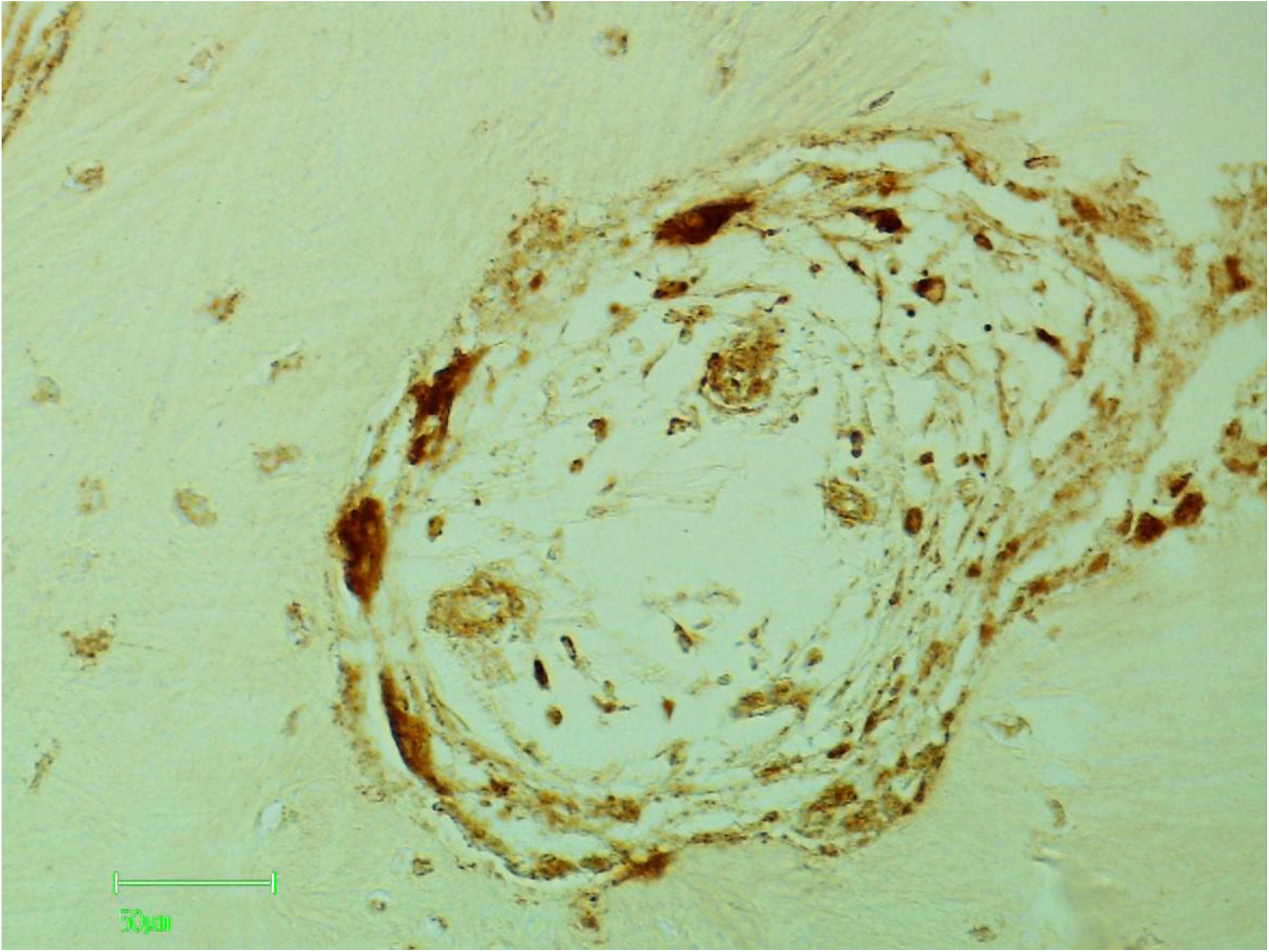
Osteoclasts, mast cells and blood vessels are stained by HHA lectin indicating the presence of N-glycans.

**Figure H.**
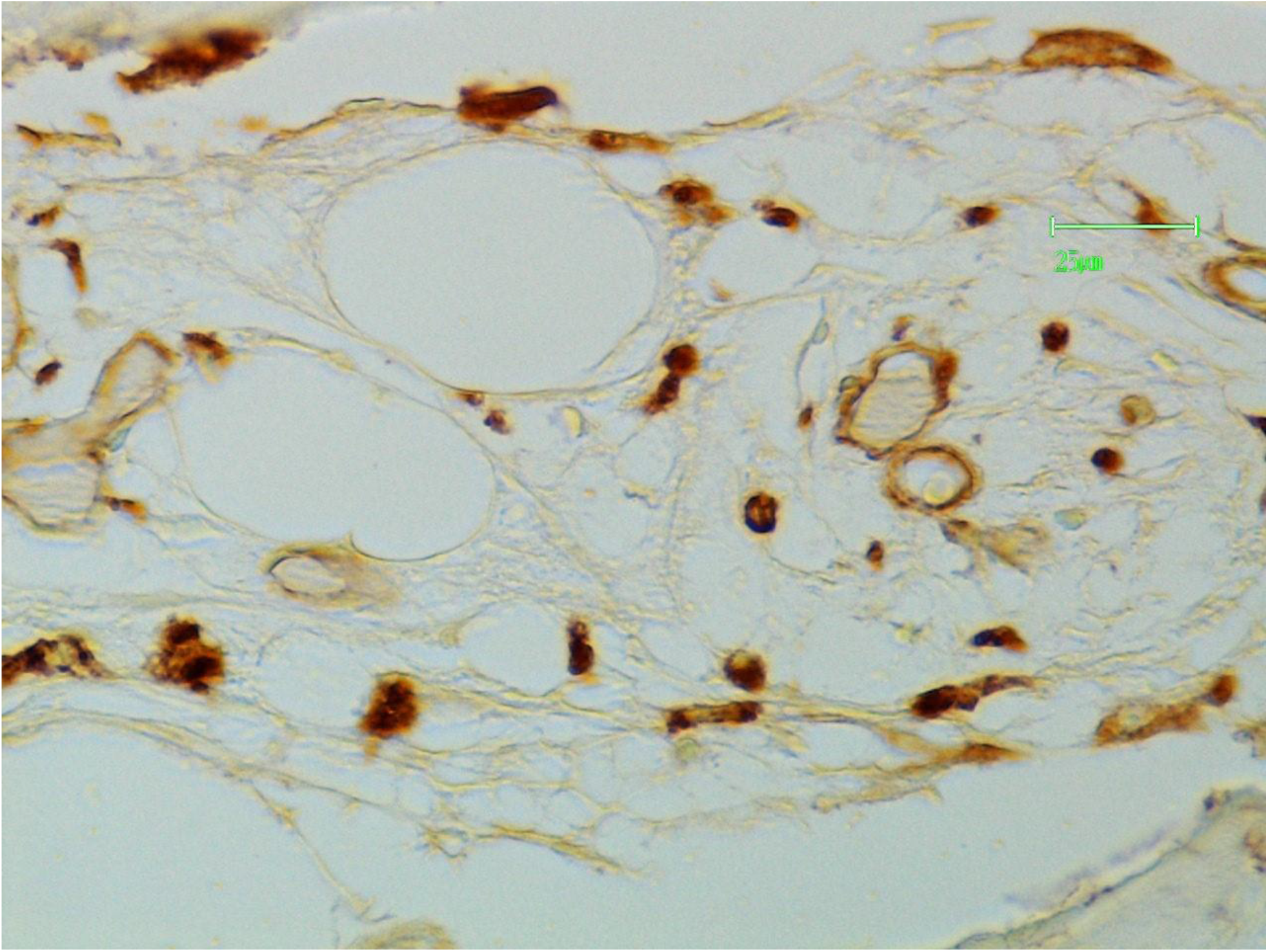
Blood vessels and mast cells are stained by lPHA lectin indicating the presence of complex N-glycans.

**Figure I.**
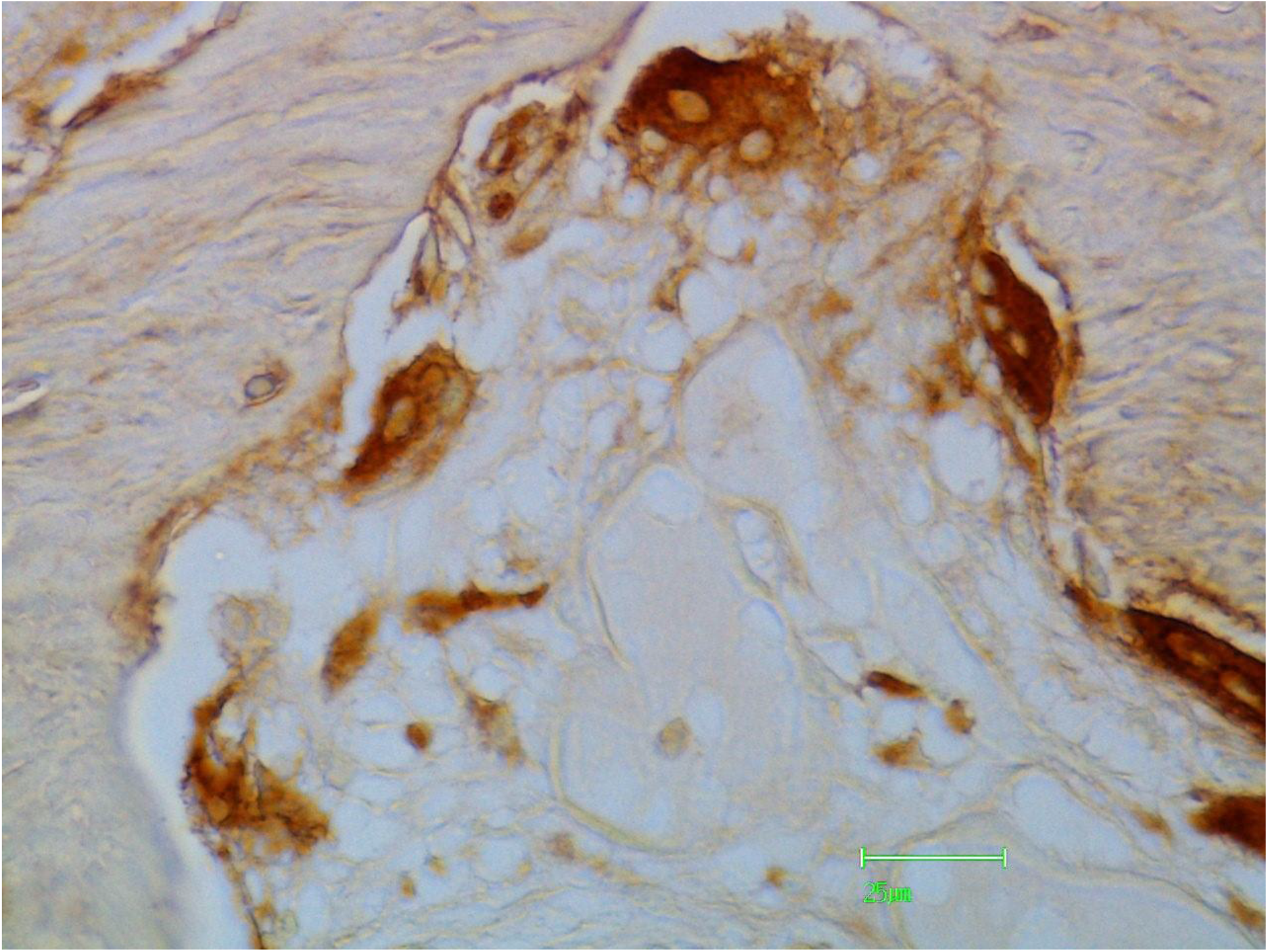
Osteoclasts eroding bone tissue are stained by sWGA lectin indicating the presence of O-glycans.

**Figure J.**
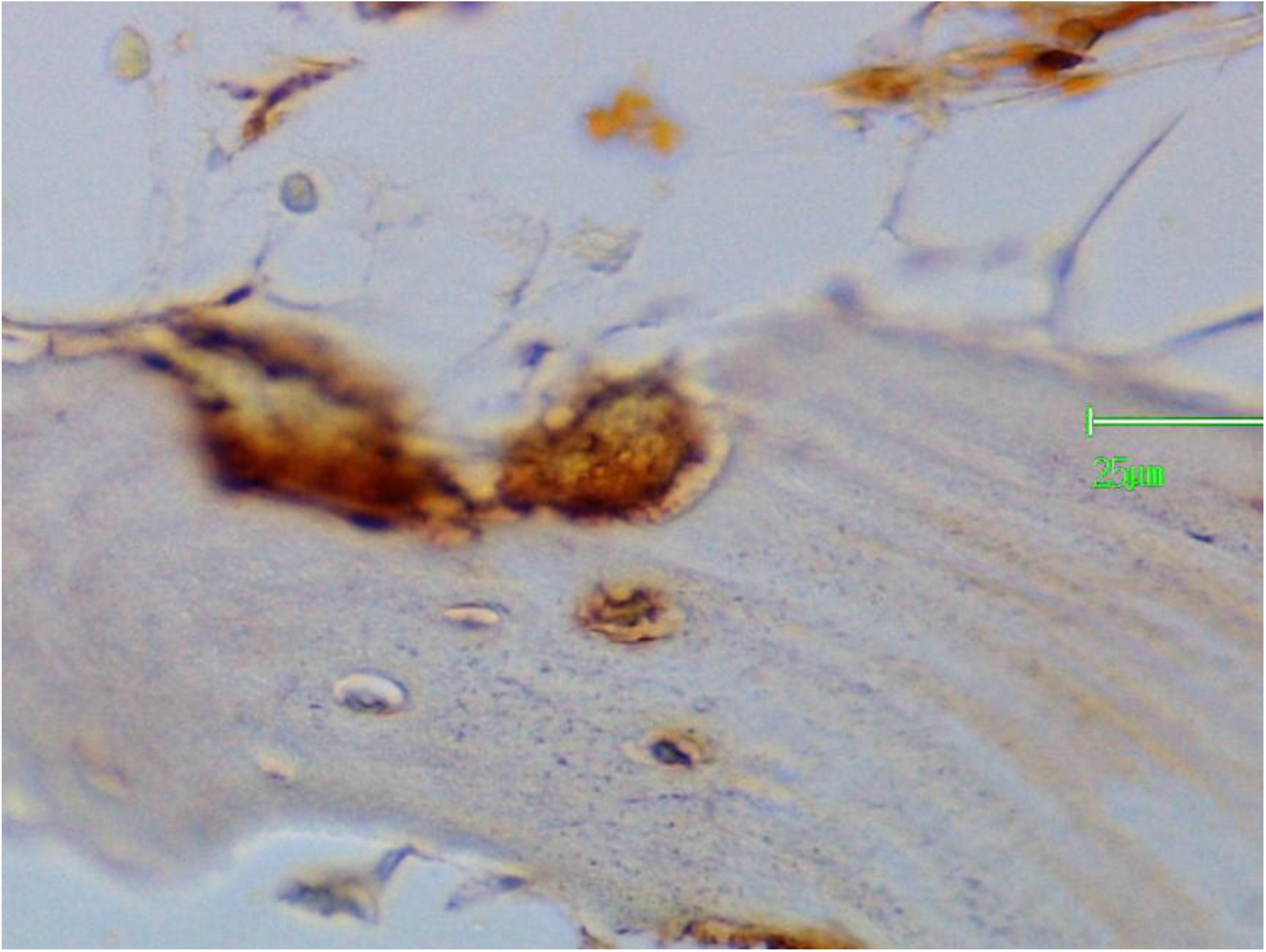
Osteoclasts are stained by lPHA lectin indicating the presence of complex N-glycans.

**Figure K.**
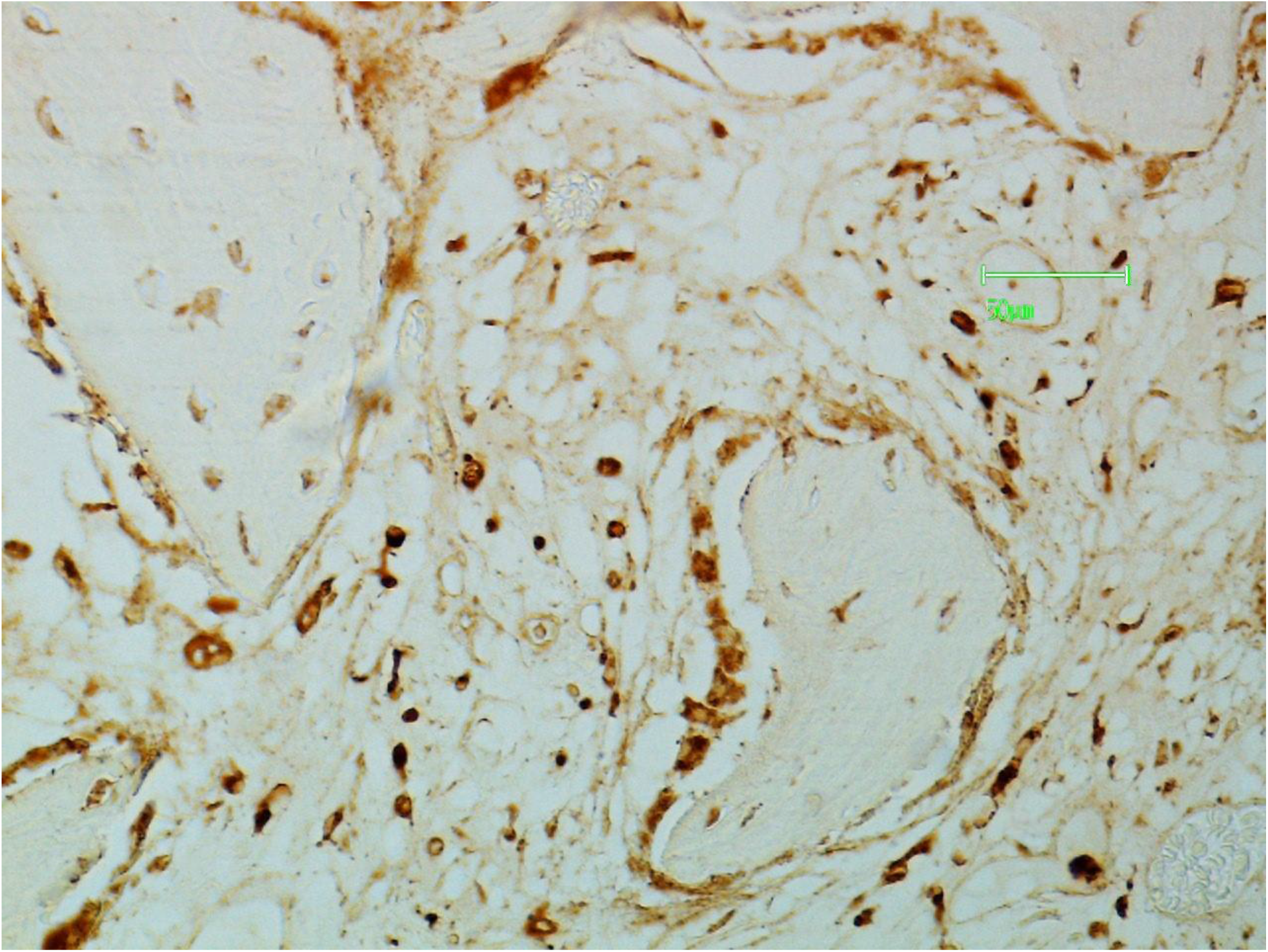
Mast cells, osteoclasts and osteoblasts are stained by HHA lectin indicating the presence of N-glycans.

**Figure L.**
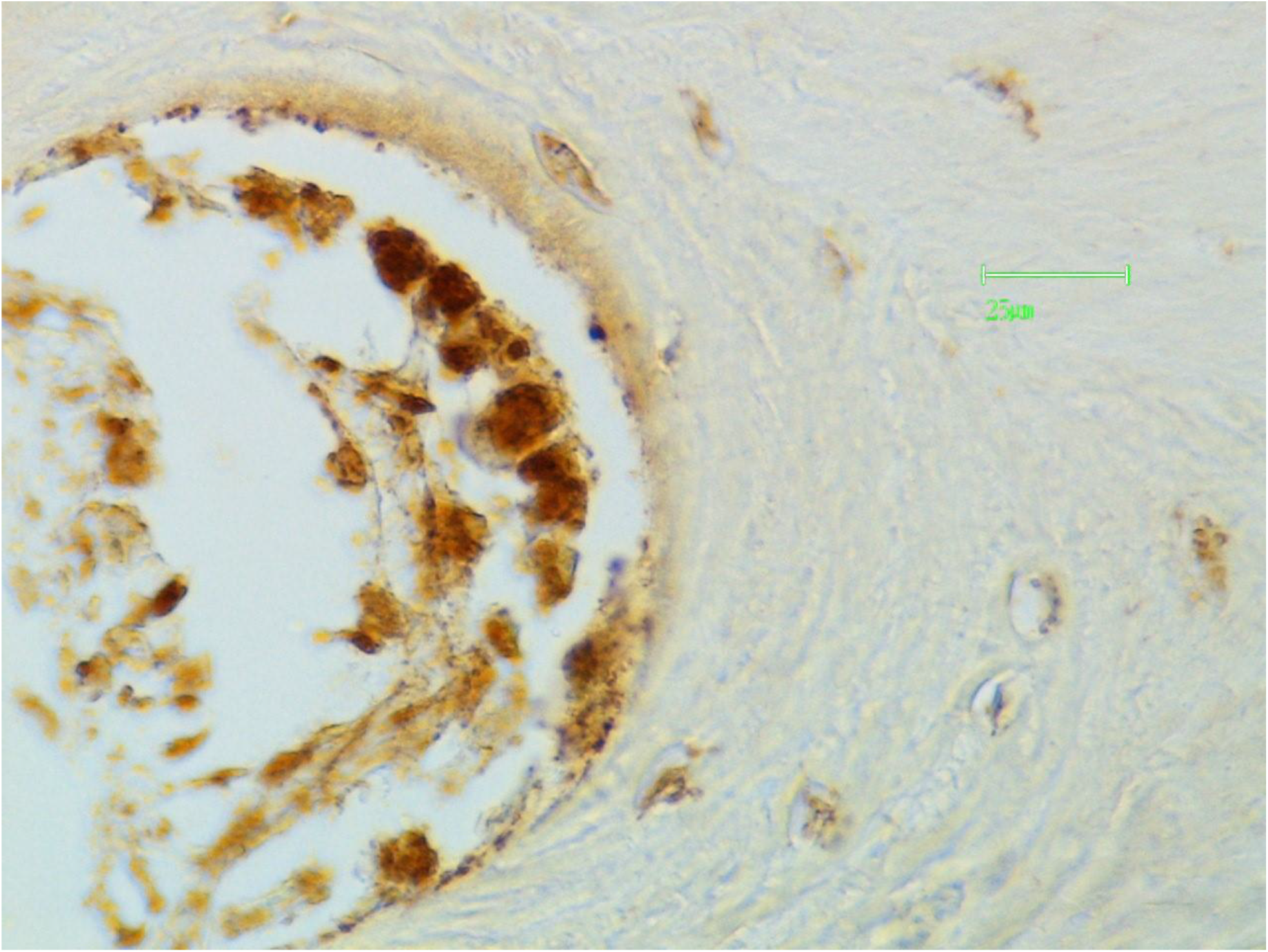
Osteoblasts are stained by HHA lectin indicating the presence of N-glycans.

**Figure M.**
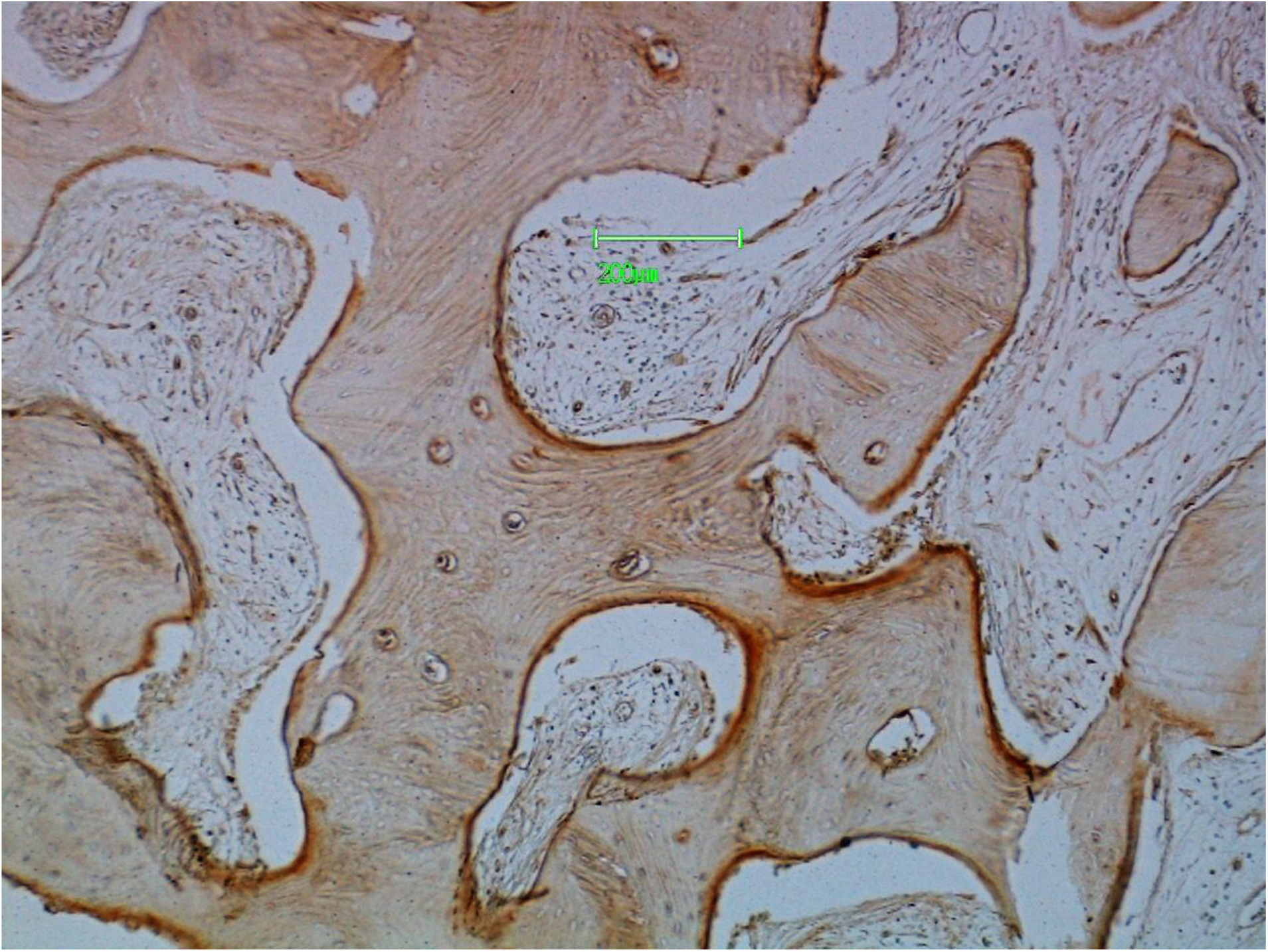
Trabecular lining cells are stained by SBA lectin indicating the presence of O-glycans.

**Figure N.**
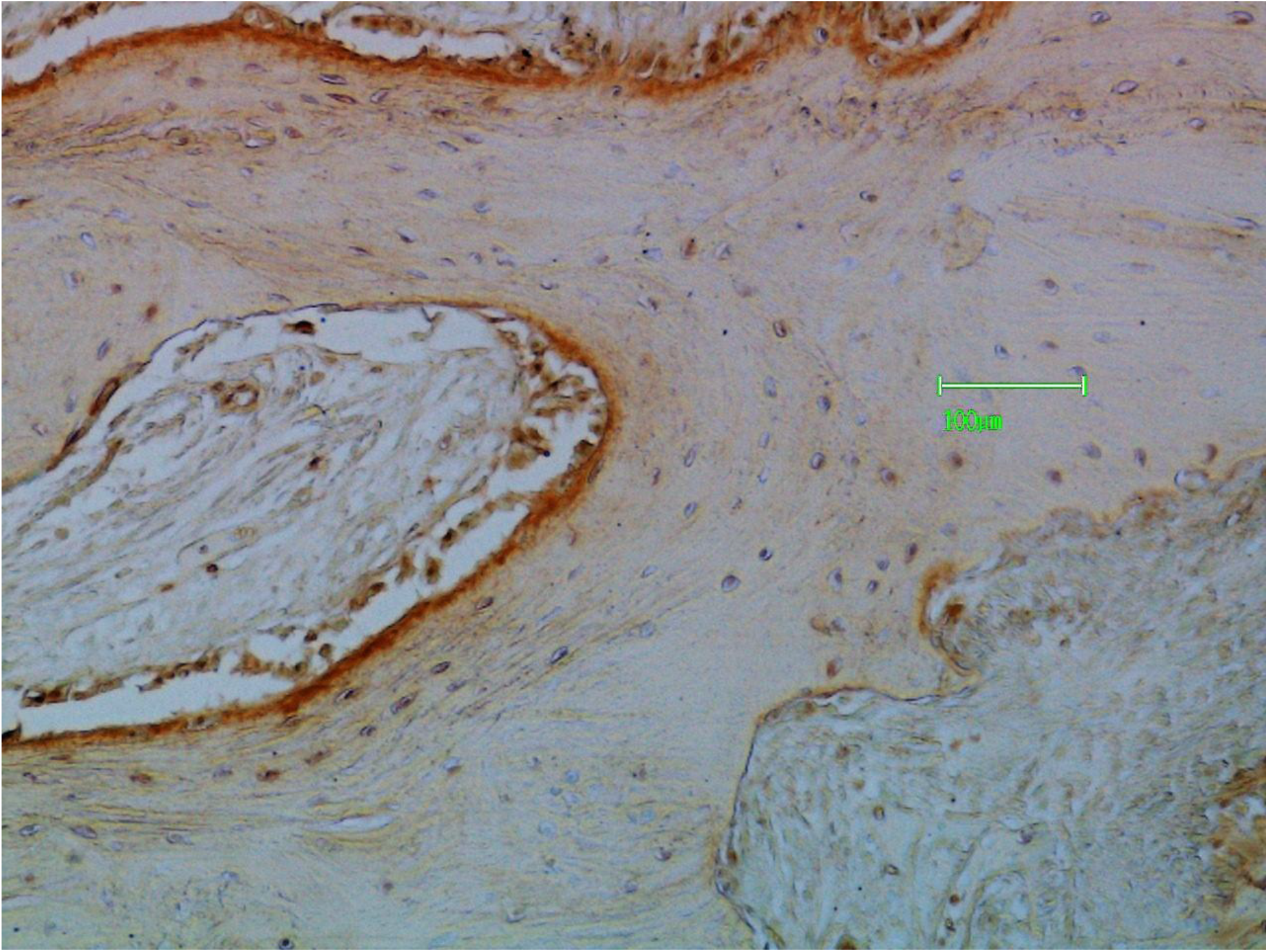
Trabecular lining cells are stained by sWGA lectin indicating the presence of O-glycans.

**Figure O.**
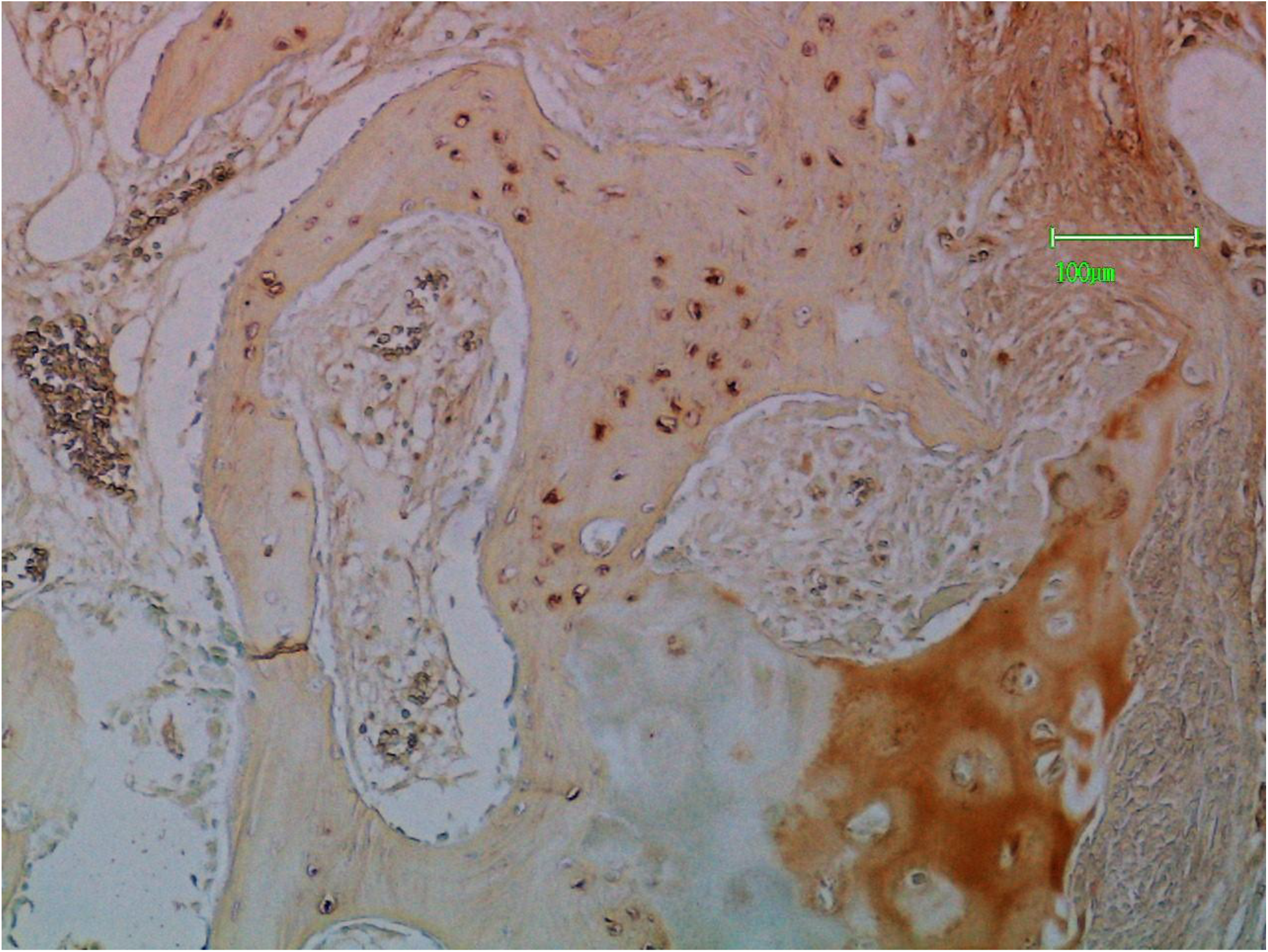
Osteocytes and the matrix of the hypertrophic cartilage zone are stained by AHA lectin indicating the presence of O-glycans.

**Figure P.**
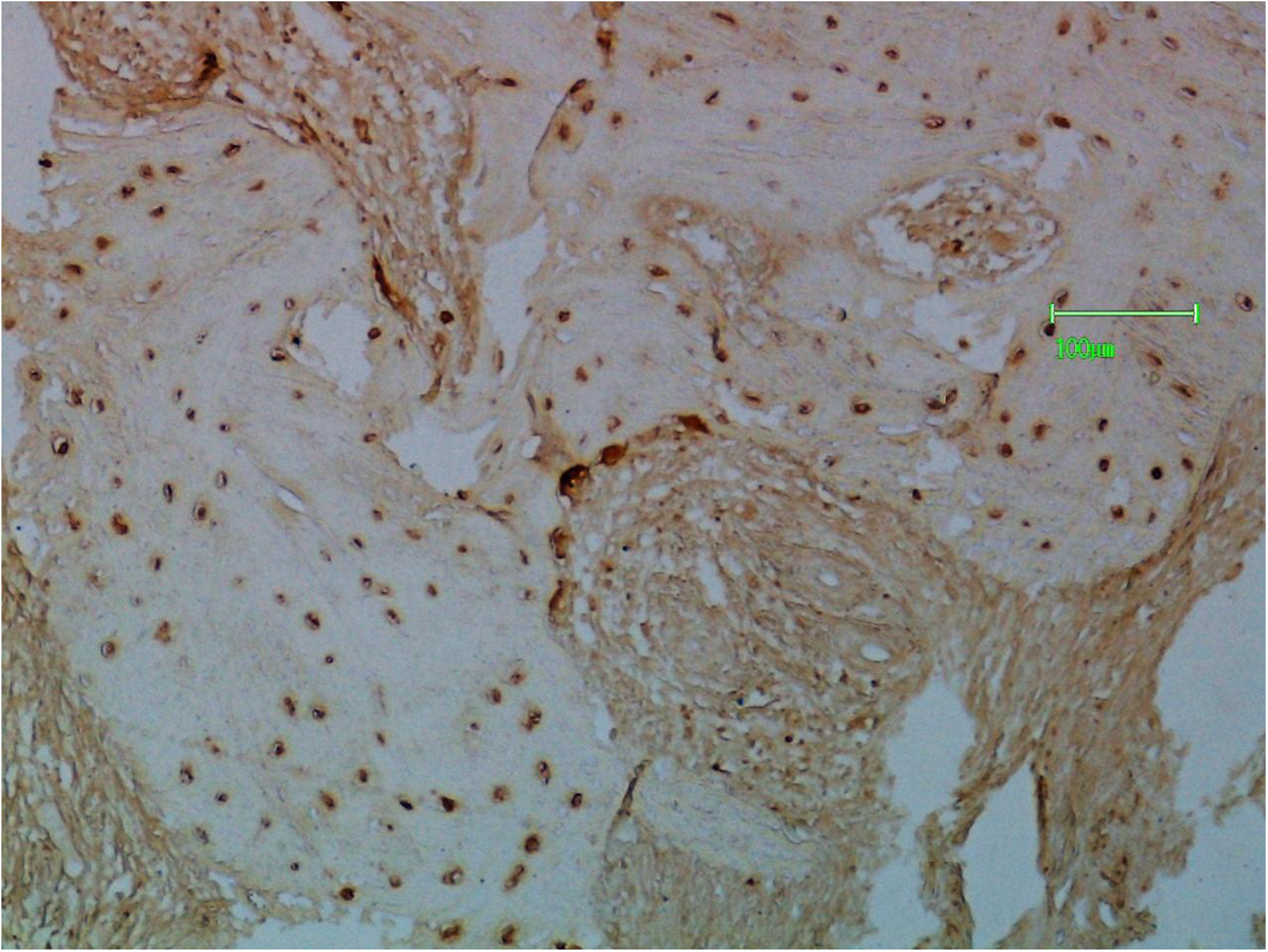
Osteocytes and osteoclasts are stained by MPA lectin indicating the presence of O-glycans.

**Figure Q.**
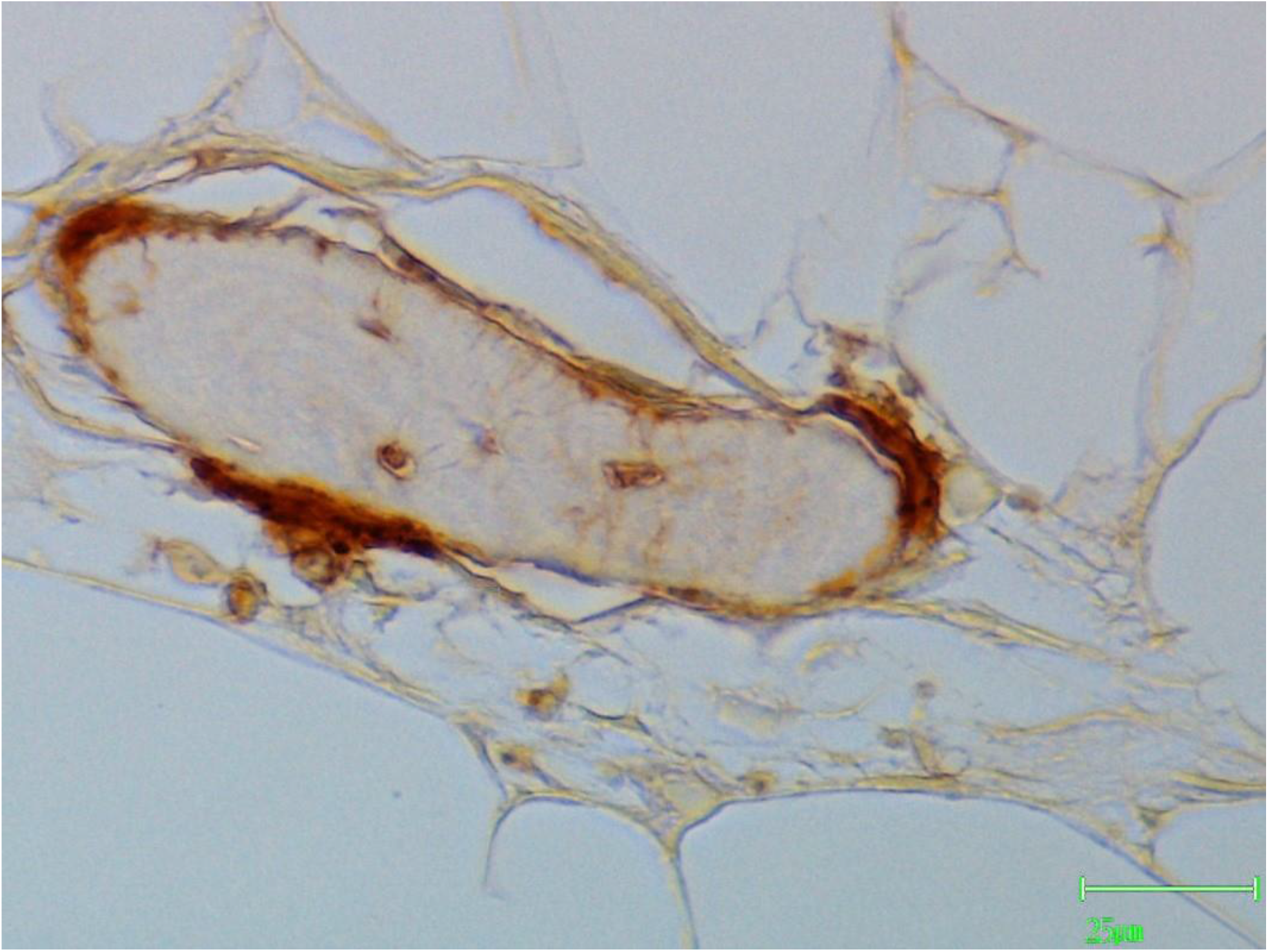
Osteocytes and trabecular lining cells are stained by MPA lectin indicating the presence of O-glycans. The intracanalicular processes are also stained.

## REFERENCES

1. Hart GW, Copeland RJ. Glycomics hits the big time. Cell 2010; 143: 672–6

2. Toegel S, Bieder D, Andre S et al. Glycophenotyping of osteoarthritic cartilage and chondrocytes by RT-qPCR, mass spectrometry, histochemistry with plant/human lectins and lectin localization with a glycoprotein. Arth Res and Ther 2013; 15: R147.

3. Lyons TJ, Stoddard RW, McClure SF, McClure J. The tidemark of the chondro -osseous junction of the normal human knee joint. J Mol Histol 2005; 36: 207–15.

4. Lyons TJ, Stoddard RW, McClure SF, McClure J. Lectin and other histochemical studies of the articular cartilage and chondro-osseous junction of the normal knee joint. J Mol Histol 2007; 38: 13–23.

5. Barkhordari A, Stoddard RW, McClure SF, McClure J. Lectin histochemistry of normal human lung. J Mol Histol 2004; 35: 147–56.

6. Stoddard RW, Jones CJ. Lectin histochemistry and cytochemistry – light microscopy : avidin-biotin amplification on resin-embedded sections. In: Rhodes JM, Milton JD (eds).Lectin methods and protocols. Humana Press 1998. Totowa NJ, pp 21–39.

7. Illes T, Fischer J. Distribution of lectin binding glycoprotein in osteoclasts. Histochemistry 1989; 91: 55–9.

8. Illes T, Fischer J, Szabo G. Lectin histochemistry of pathological bones. Bull Hosp Jt Dis 1999; 58: 206–11.

9. Kirkpatrick CJ, Jones CJ, Stoddard RW. Lectin histochemistry of the mast cell: a light microscopical study. Histochem J. 1988; 20: 139–46.

10. Herzog BH, Fu J, Xia L. Mucin-type O-glycosylation is critical for vascular integrity. Glycobiology 2014; 24: 1233–41.

11. Dennis JW, Laferte S, Waghorne C, Britman ML, Kerbel RS. Beta 1-6 branching of Asn-oligosaccharide is directly associated with metastasis. Science 1987; 236: 582–5.

12. Tian E, Ten Hagen KG. Recent insights into the biological roles of mucin-type O-glycosylation. Glyconj J 2009; 26: 325–34.

13. McClure J, Smith PS. Oncogenic osteomalacia. J Clin Pathol 1987; 40: 446–53.

14. Wilson KM, Jagger AM, Walker M et al. Glycans modify stem cell differentiation to impact on the function of resulting osteoclasts. J Cell Sci. 2018 Feb 15; 13(4):jcsz09452. DoI 10.1242/jcs.209452.

15. Guy Y-C, Yuan Q. Fibroblast growth factor 23 and bone mineralization. Int. J Oral Sci 2015; 7: 8–13.

16. Yamasita T, Yoshioka M, Itoh N. Identification of a novel fibroblast growth factor FGF-23, preferentially expressed in the ventrolateral thalamic nucleus of the brain. Biochem Biophys Res Commun 2000; 277: 949–8.

17. Bonewald LF, Walker MJ. FGF-23 production by osteocytes. Pediatr Nephrol 2013; 28: 563–8.

18. Wang H, Yosiko Y, Yamamoto R et al. Overexpression of fibroblast growth factor 23 suppresses osteoblast differentiation and matrix mineralization in vitro. J Bone Miner Res 2008; 23: 939–48.

19. Sitara D, Kim S, Rizzique MS et al. Genetic evidence of serum phosphate-independent functions of FGF23 on bone. PLoS Genet 2008; 4: 1–10.

20. Shaloub V, Ward SC, Sun B. Fibroblast growth factor 23 (FGF23) and alpha Klotho stimulate osteoplastic MC3T3. E1 cell proliferation and inhibit mineralization. Calcif Tissue Int 2011; 89: 140–50.

21. Bonewald LF. The amazing osteocyte. J Bone Min Res 2011; 26: 229–38.

22. Bullough PG, Jagannath A. The morphology of the calcification front in articular cartilage. Its significance in joint function. J Bone Jt Surg B 1983; 65: 72–8.

23. Gannon JM, Walker G, Fischer M et al. Localization of type X collagen in canine growth plate and adult articular cartilage. J Orthop Res 1991; 9: 485–94.

24. McClure SF, Stoddard RW, McClure J. A comparative study of lectin binding to cultured chick sternal chondrocytes and intact chick sternum. Glycoconj J 1997; 14: 365–77.

25. Li RC, Wong MY, DiChiara AS et al. Collagen’s enigmatic, highly conserved N-glycan has an essential proteostatic function. PNAS 2021; 118: e2026608118.

26. Chambers TJ. The regulation of osteoclastic development and function. Cell and molecular biology of vertebrate hard tissue 2007. Ciba Foundation Symposium 136: 92-107. Eds: Evered D, Harnett S.

27. Fau X, Wu X, Crawford R et al. Macro, micro and molecular changes of the osteochondral interface in osteoarthritis development. Frontiers in Cell and Developmental Biology 2021; 9: 1–17.

